# Investigating curvature sensing by the Nt17 domain of Huntingtin protein

**DOI:** 10.1101/2024.10.20.619314

**Authors:** Neha Nanajkar, Abhilash Sahoo, Shelli L. Frey, Silvina Matysiak

## Abstract

Nt17, the N-terminal domain of the huntingtin protein (htt), has garnered significant attention for its role in htt’s membrane binding and aggregation processes. Previous studies have identified a nuclear export sequence within the Nt17 domain and demonstrated its localization at various cellular organelles. Recent evidence suggests that, like other amphipathic helices, Nt17 can sense and preferentially bind to curved membranes. Gaining deeper insight into this behavior is essential to fully understand the function of this domain.

In this study, we combine coarse-grained molecular dynamics simulations with circular dichroism (CD) spectroscopy to investigate the mechanism behind Nt17’s curvature sensing. We generated a unique hemispherical-planar membrane model, where 36% of the upper leaflet surface is curved, allowing us to evaluate Nt17’s binding preferences. Our findings show that Nt17 exhibits a strong preference for curved regions, with approximately 78 ± 7 % of peptides binding to these areas. This interaction is primarily mediated by the terminal Phe residues, indicating that Nt17’s curvature sensing is driven by its ability to detect lipid packing defects. Furthermore, Nt17 not only senses these defects but also amplifies them by coalescing smaller pockets. Mutating the Phe residues to methionine, a smaller hydrophobic residue, significantly reduces Nt17’s curvature sensitivity, resulting in equal binding to both curved and planar regions. CD spectroscopy corroborates these results, showing that Nt17 binds more strongly to highly curved small unilamellar vesicles (SUVs) compared to larger, less curved large unilamellar vesicles (LUVs).

## Introduction

Huntingtin (htt) is an intracellular protein implicated in the development of Huntington’s disease (HD). In HD, htt has an aberrant expansion of the polyglutamine region, resulting in the formation of toxic aggregates, subsequently leading to neurodegeneration. ^1^ The flanking domains of polyglutamine, Nt17 (MATLEKLMKAFESLKSF) and proline rich domains are known to promote and impede aggregation respectively.^2^ Nt17, referring to the first 17 residues in the N-terminus, is particularly important with respect to membrane interaction. Composed of hydrophobic and polar residues, this fragment is disordered in solution but forms an amphipathic α-helix when in contact with membranes or when the protein is in aggregated form.^3,4^ Nt17’s ability to self-associate results in higher rates of htt aggregation and fibril formation through an increase in local density of the polyglutamine stretch.^2,5^ It also plays a pivotal role in cellular functioning. Studies indicate that Nt17 acts as the membrane anchoring domain of the first exon of htt, with reports of subcellular localization at the mitochondria, endoplasmic reticulum and the Golgi apparatus.^6–8^ A leucine rich nuclear export motif present within the fragment is responsible for the localization of the htt protein.^5^

Specific mutations or complete deletion can result in nuclear accumulation of htt and a subsequent rise in cell toxicity. Nt17 is also the target of post-translational modifications that alter its aggregation and localization propensities. ^9,10^ Phosphomimetic mutations to S13/S16 of the Nt17 domain resulted in reduced cellular toxicity even for mutant htt fragments having extended polyQ domains.^11^ Targeted modifications of the Nt17 domain are being considered a possible avenue for the treatment of HD.^12^ These and other works underscore the significance of the Nt17 domain in Huntington’s Disease pathology.

Recent reports have suggested that Nt17 has a preference towards curved membranes. Studies using EPR spectroscopy demonstrate that the percentage of Nt17 binding to vesicles reduces as vesicle diameter increases.^13^ In addition, curvature affinity of the synthetic N-terminal htt peptide mimetic (Nt17Q35P10KK) was observed through atomic force microscopy.^14^ Silica nanoparticles were placed under a lipid bilayer to create regions of artificial curvature of approximately 50 nm in diameter. Upon incubating the bilayer with Nt17Q35P10KK, the peptides accumulated on the curved rather than planar regions of the bilayer, indicative of preferential binding.

Curvature sensing has been observed in other peptides associated with neurodegeneration, such as α-synuclein and amyloid-*,8*,^15,16^ suggesting a relationship between neurodegeneration and membrane curvature. α-synuclein and amyloid-*,8* are known to interact with membranes in ways that may disrupt cellular processes. These peptides often accumulate at sites of high membrane curvature, which can destabilize membrane integrity or induce abnormal curvature.^17–19^ This may lead to altered trafficking, vesicle formation, or even membrane rupture. Over time, such disruptions could contribute to the toxic aggregation of proteins and mitochondrial dysfunction, all hallmark features of neurodegeneration. Prior work on curvature sensing has attributed sensing to electrostatic interactions or the hydrophobic effect. *ff*synuclein and ALPS, two commonly studied curvature sensors, were found to have differing modes of curvature sensing.^20^ α-synuclein was found to have a poorly developed hydrophobic face, resulting in its sensitivity to anionic curvatures. In contrast, ALPS motifs have a very well developed hydrophobic face and few charged residues, leading to a hydrophobic effect driven sensing mechanism. It is unclear which of these categories Nt17 falls into, necessitating further examination.

Experimental and computational studies performed so far have unveiled a great deal about the interactions of Nt17 with lipids. All atom molecular dynamics simulations revealed characteristic features of Nt17 binding to zwitterionic lipid membranes,^21,22^ including tilt angle, insertion depth, and the effects of binding on the bilayer itself. Experimental studies using solid-state NMR and circular dichroism have provided valuable insights into Nt17’s structure in both solution and membrane-bound states. These experiments revealed significant changes in membrane thickness, lipid order parameters, and the structural transitions of Nt17, such as the shift from a random coil to an alpha-helix, which was shown to be dependent on the lipid environment. This work has been essential in enhancing our understanding of how Nt17 interacts with membranes in different contexts. However, molecular level insights into the mechanisms responsible for curvature sensing are still missing. Traditional all atom MD is capable of modeling systems in high resolution but, it is computational challenging to capture large-scale biological phenomena such as membrane binding and peptide aggregation. Coarse-grained molecular dynamics (CG-MD) simulations, however, are uniquely poised to address these shortcomings. In CG-MD, similar atoms are grouped together to form beads. By “coarse-graining” the system, larger systems can be simulated, and for longer periods of time.

In this work, CG-MD is applied to simulate the binding of Nt17 to a membrane architecture containing both curved and planar regions, as depicted in Figure 1. As expected, we find that Nt17 does in fact sense curvature, with the bulk of binding occuring at regions with lipid packing defects. The terminal Phe residue drives the interaction process, with peptide adsorption only occurring after partitioning of the two Phe residues (F11 and F17). Mutation of the Phe residues to M, a smaller hydrophobic amino acid, suppresses the curvature sensing ability. Circular dichroism experiments performed on Nt17 and its mutant, Nt17*^F^* ^11*M/F*^ ^17*M*^, are in agreement with these findings. Finally, we show that Nt17 seeks out regions with higher area per lipid, such that the bulky sidechains of Phe can partition to the membrane, at regions of lipid packing defects. Since Nt17*^F^* ^11*M/F*^ ^17*M*^ has smaller sidechain residues it is capable of partitioning to regions without packing defects.

**Figure 1:**
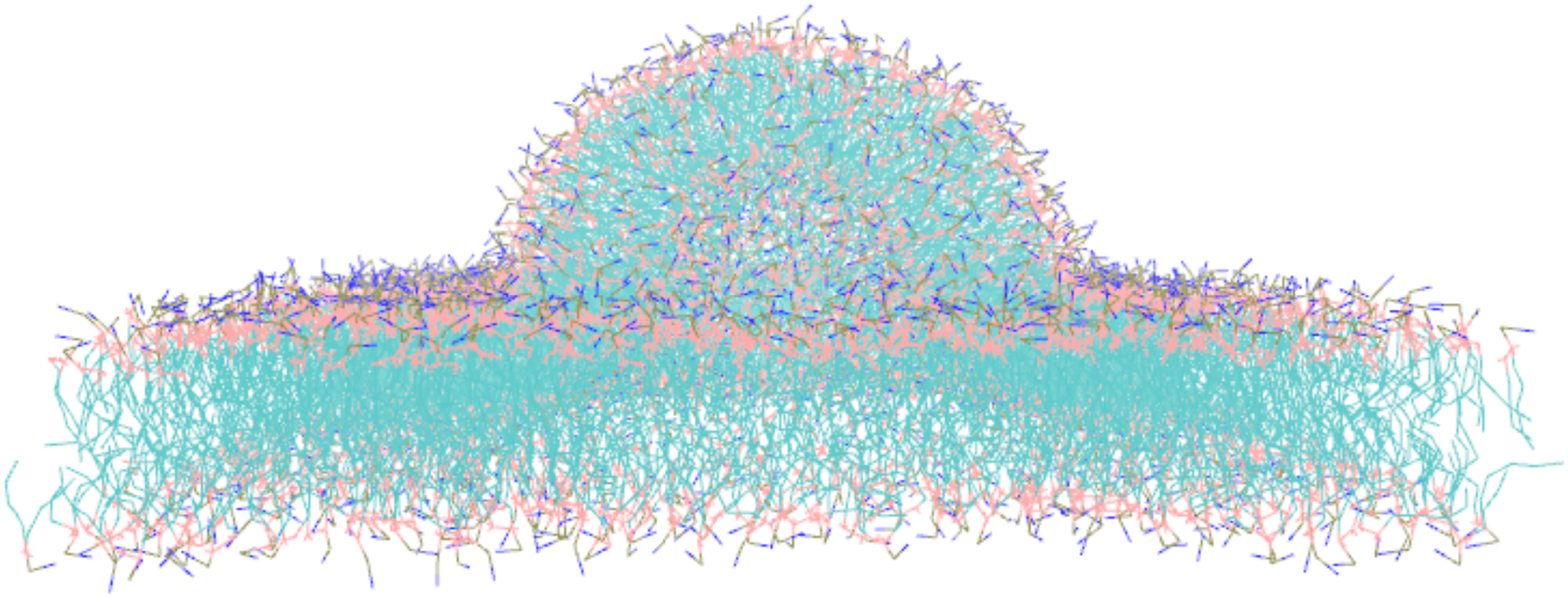
Hemispherical-planar membrane architecture used in this study, consisting of both planar and curved regions. The curved region of the upper leaflet accounts for 36% of the upper membrane surface area.

## Results and Discussion

### Nt17 prefers interacting with curved regions of the membrane

To assess whether Nt17 can sense curvature, we ran simulations of Nt17 with a hemispherical-planar membrane architecture, pictured in Fig. 1. The curved region of the membrane (M*_c_*) accounts for around 36% of the total surface area, while the planar region (M*_p_*) makes up the remaining 64%. Thus, to characterize Nt17 as a curvature sensor, an excess of 36% of the peptides in the system must bind to the curved region. Fig. 2A presents a timeseries analysis of percentage of Nt17 binding for one replicate, either interacting with M*_c_*, M*_p_*.

**Figure 2:**
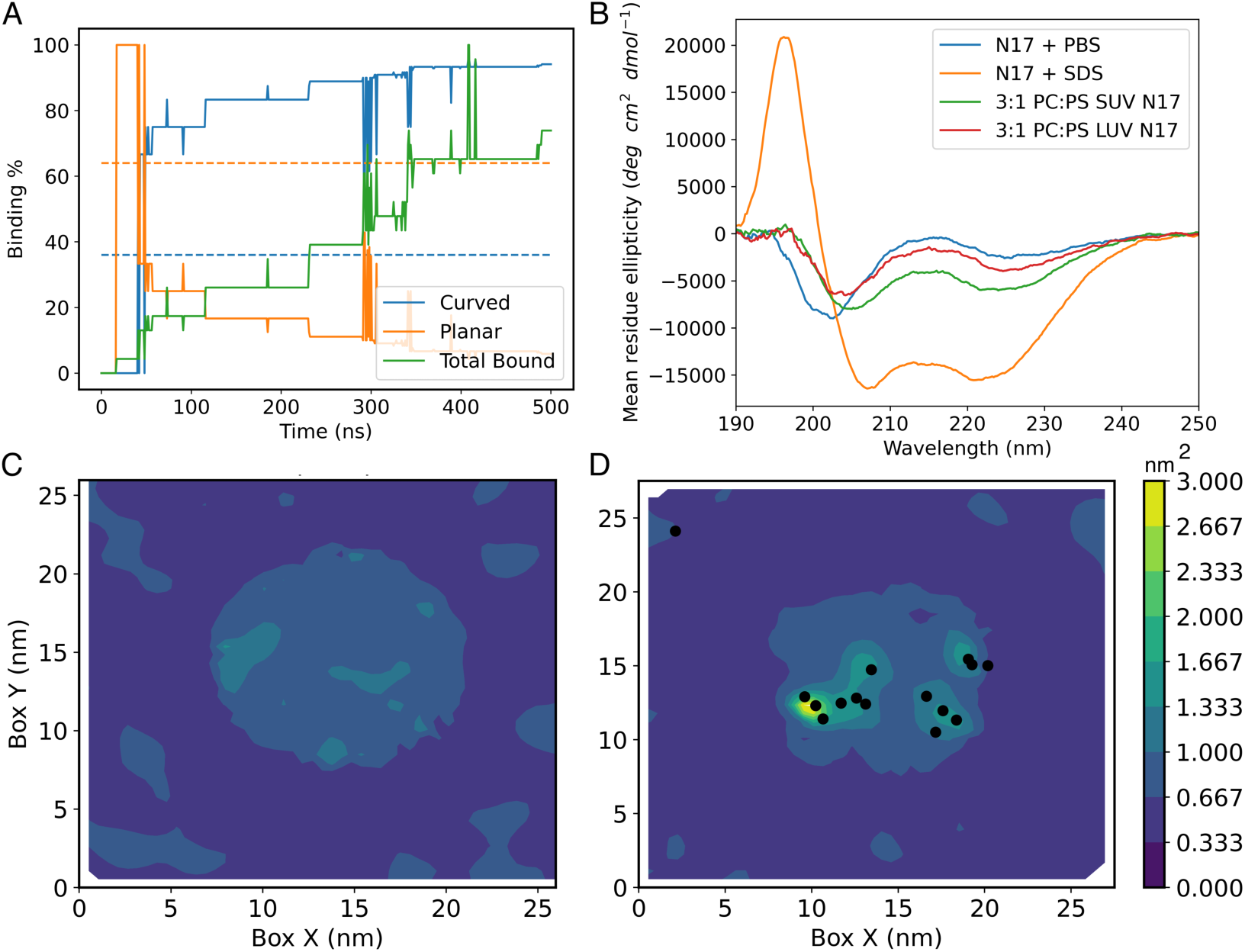
Nt17 preferentially binds to curved regions of the membrane. A) Binding percentages for one replicate of Nt17 to the curved and planar regions are denoted blue and purple respectively. The dotted lines represent the minimum percent binding required to qualify as having preference for curved and planar regions. The solid green line represents the percentage of membrane bound peptides in the system, B) Circular dichroism spectra for Nt17 in varying solutions. Circular dichroism of Nt17 in varying solutions. 10 mM pH 7.3 PBS (blue); 10 mM pH 7.3 PBS with 2 mM sodium dodecyl sulfate (SDS) to induce helical folding (orange); 3:1 POPC:POPS SUVs in 10 mM pH 7.3 PBS (green); and 3:1 POPC:POPS LUVs in 10 mM pH 7.3 PBS (red). SUVs (small unilamellar vesicles) are around 30nm in diameter and LUVs (large unilamellar vesicles) are around 120 nm in diameter, C) Area per lipid (APL) plot of the control simulation (no peptides), D) APL plot for one replicate of Nt17. Black dots indicate the center of mass of peptides that have bound to the membrane.

To calculate the binding percentage, we summed up the peptides directly and indirectly bound to specific regions of the membrane (M*_c_* or M*_p_*) and normalized by the number of peptides interacting with the membrane at that time point. In the first 50ns of the simulation, the planar binding percentage is high, but this can be explained but the low overall binding, denoted in green. The percentage of peptides bound to M*_c_* increases to around 85% (blue) by the end of the simulation, far outweighing the 36% required to qualify Nt17 as a curvature sensor. Other replicates for Nt17 follow a similar trend (Figure S1), clearly positioning Nt17 as a curvature sensor. Note, at the studied timescales, we do not see complete binding of Nt17 to the membrane surface. Instead, Nt17 forms aggregates in solution that are, on average, 8-10 peptides in size (see Figure S2A).

Experimental verification of these findings was carried out by collecting circular dichroism (CD) spectra on Nt17 in the presence of small and large unilamellar vesicles. Small unilamellar vesicles (SUVs) have a diameter of approximately 30 nm and the large unilamellar vesicles (LUVs) have a diameter of approximately 120 nm. As vesicle size increases, its curvature reduces. Thus, a curvature sensing peptide should then prefer binding to an SUV over an LUV. To enhance the CD signal in peptide binding experiments, the addition of anionic lipids is required, because the membrane partitioning constant of Nt17 with SUVs is 2 orders of magnitude larger in 3:1 POPC:POPS mixtures when compared to 100% POPC.^22^ Of note, both NMR and atomistic simulations have shown Nt17 binding and helix formation in 100% POPC bilayers,^21–23^ as observed in our simulations. We observed a tilt angle of 99.3 ± 7.9 for single Nt17 peptide interacting with the membrane, in good agreement with prior works.^23^ Figure 2B shows the spectra for Nt17. In buffer, Nt17 adopts a random coil structure, while in 2 mM SDS Nt17 is alpha helical, with defined minimas at 208 and 222 nm. When bound to the vesicles, Nt17 adopts a combination of random coil and helical structures, evident through the minimas at 205 and 222 nm. However, in the presence of SUVs (green), Nt17 exhibits a significantly more intense spectra, indicating more alpha helical signal and more binding overall.

Figure 2 C, D present overhead views of area per lipid (APL) for the hemispherical-planar membrane architecture simulations in control and with peptides respectively. In the planar region L*_d_* is predictably lower, as lipids tend to be more ordered and closely packed. There is a marked increase in APL at the curved region because of the formation of lipid packing defects. Figure 2D presents the APL at the last frame of one 500 ns replicate. Here, the black dots represent the center of mass of the peptides interacting with the membrane. For this replica, the majority of Nt17 binding occurs at M*_c_*, with only one peptide binding to M*_p_*. The APL plots for the 2 other replicates reflect this trend, with most binding occurring at M*_c_* (see Figure S3) Thus, we confirm that Nt17 is sensitive to and prefers binding to curved membranes.

### Bulky hydrophobic residues are necessary for sensing curvature

We calculated the probability of first membrane interaction across all residues of Nt17 to better understand how Nt17 interacts with the membrane. Fig. 3A reveals that about 30% of the initial peptide-lipid interaction occurs through the terminal Phe residue, highlighting the importance of large hydrophobic groups in the curvature sensing process. In addition, there is a 20% chance the peptide interacts with the membrane via the first four residues located on the N-terminus (M through L). Nt17 forms small aggregates in solution prior to interacting with the membrane, and the peptides orient their hydrophobic sides facing inward, forming helical bundles, resulting in residues at either termini being more likely to participate in the first interaction. Bundling of Nt17 has been proposed in a number of prior studies.^24–26^ Figure S2A presents the average aggregate size for Nt17 interacting with M*_c_*, M*_p_* and remaining in solution. The average size of aggregates on M*_p_* and M*_c_* are 3 and 5 respectively. Because Phe is significantly more hydrophobic than MET, thus it is expected that the first interaction would occur primarily through the terminal Phe residue.

**Figure 3:**
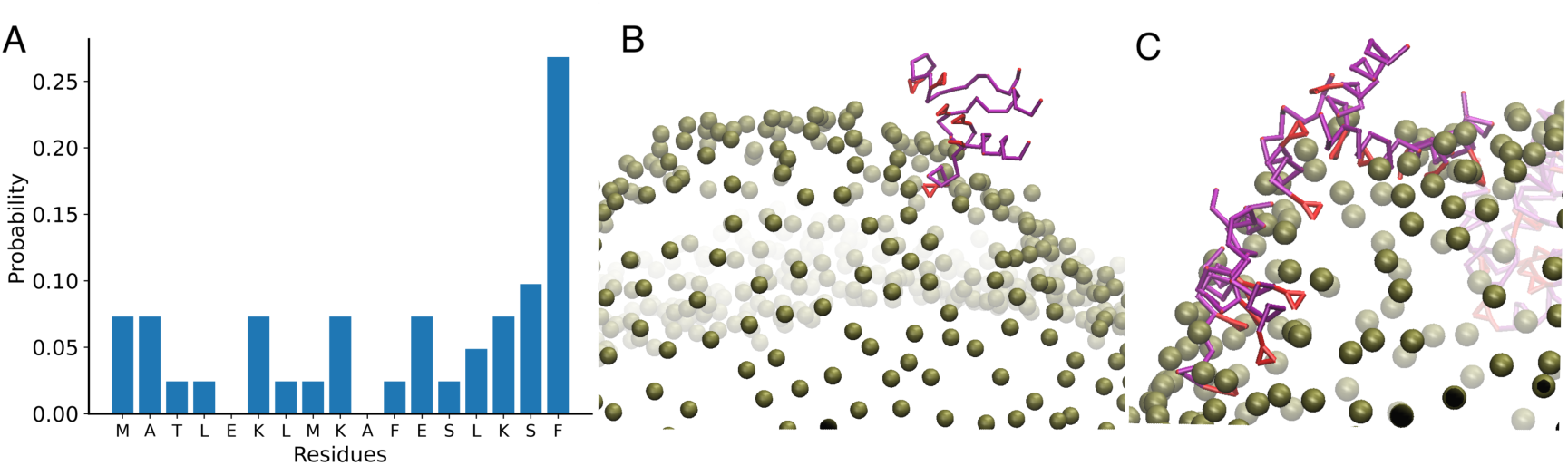
Nt17-membrane interaction greatly depends on the Phe residues. A) The bulk of Nt17-membrane interactions occur through the C-terminal Phe residue, B) Snapshots of Nt17 interacting with the curved region of the upper leaflet of the lipid membrane and C) Nt17 partitioning to the upper leaflet upon insertion of the Phe residues. Protein backbones and Phe sidechains are denoted in purple and red respectively. Phosphate beads of the lipid are colored in olive green.

Figs. 3B and C present snapshots of the peptide interacting with the membrane. Nt17 first approaches the membrane, followed by interaction of the terminal Phe residue with the membrane surface. Only when both F11 and F17 residue insert to the surface does Nt17 partition completely to the membrane. In Fig. 3C, we note multiple peptides adsorbed to the curved region of the membrane, with deep insertion of the Phe residues (red triangles).

### Removal of Phe residues leads to a loss in curvature sensitivity

Insertions of hydrophobic residues into regions with lipid packing defects is one method by which amphipathic helices can sense, and in some cases generate curvature. In particular, the ALPS motif senses deep lipid packing defects through the insertion of large hydrophobic residues.^27^ Having established the role of the Phe residues in membrane interaction, we tested a mutated version of Nt17, henceforth called Nt17*^F^* ^11*M/F*^ ^17*M*^, where the bulky Phe residues in positions 11 and 17 were mutated to methionine. Methionine is also a hydrophobic amino acid but is smaller than phenylalanine, so the amphipathic nature of Nt17 is not disrupted. Furthermore, methionine is a residue commonly found in helical structures, thus the helical nature of Nt17 should not be perturbed.^28^ This mutation allows us test the contribution of the bulky hydrophobic groups in the curvature sensing process.

Although Figure 3A indicates that F17 is the responsible for the bulk of first interactions with the membrane, we elected to mutate both F11 and F17 to methionine. This was done to eradicate the possibility of bundling between peptides. Prior work by our group has demonstrated that Nt17 domains of htt have a propensity to bundle together, arranging so the hydrophobic residues point inwards forming a hydrophobic core.^24^ The large hydrophobic residues on Nt17 give way to two distinct and competing behaviors: either Nt17 bundles together, or it partitions to the membrane by inserting the Phe residues. Figure S4A presents a boxplot of the number of hydrophobic interactions within aggregates (in solution) of Nt17 and Nt17*^F^* ^11*M/F*^ ^17*M*^ respectively. Here, higher values indicate more interactions between the side chain beads of hydrophobic residues, ie. a larger hydrophobic core. Thus, it is clear the aggregates of Nt17 in solution experience more hydrophobic interactions. As a result of this, we see differences in average aggregate size. Where Nt17 can form aggregates in the range of 8-10 peptides, aggregates of Nt17*^F^* ^11*M/F*^ ^17*M*^ are smaller at 4-6 peptides on average, as seen in Figure S2B. Although the formation of large oligomers of Nt17 is atypical in experimental studies, simulations usually require concentrations higher than physiological concentrations to accelerate the aggregation process and to observe aggregation in a reasonable simulation time.^29^ The concentration of Nt17 in CD experiments amounts to 76 µM, while the peptide concentration in simulation is closer to 10mM. At these elevated concentrations, Nt17 aggregation is not an unexpected phenomena. Figure S4B presents a binary classification of Nt17 behavior. 60.9% of Nt17 bundle together via interpeptide Phe-Phe interactions and do not interact with the lipid membrane. Thus, by mutating both F to M, we eradicate the possibility of bundling and achieve comparable binding fractions for both the wild-type Nt17 or its mutated counterpart (Figure S4C).

Fig. 4A presents the binding percentage plot of one replicate of Nt17*^F^* ^11*M/F*^ ^17*M*^, where a loss in curvature specificity can be noted for the mutated peptide. By the end of the simulation, we see that the majority of peptide is bound to the planar region, though the percentage fluctuates around 64%. Interestingly, in the second replicate of Nt17*^F^* ^11*M/F*^ ^17*M*^, we see no binding to the curved region. The third replicate starts off with high planar binding percentage, but at the 400ns mark, the curved binding percentage overtakes it. Visualization of this simulation revealed that peptides that had initially bound to the planar region diffused to the curved and vice versa, resulting in the fluctuating binding percentages. Thus, we can conclude that curvature sensing is drastically reduced with Nt17*^F^* ^11*M/F*^ ^17*M*^. With Nt17, all replicas had a clear curvature preference, but for Nt17*^F^* ^11*M/F*^ ^17*M*^ each run varies. The circular dichroism spectra of Nt17*^F^* ^11*M/F*^ ^17*M*^, shown in Fig. 4B, is markedly different than that of wild-type Nt17. Nt17 demonstrates a clear preference towards SUVs (more curvature) over LUVs, demarcated by the significant increase in spectral intensity (Fig. 2B). On the other hand, Nt17*^F^* ^11*M/F*^ ^17*M*^ only indicates a slight preference for SUVs over LUVs. This indicates that removal of the bulky hydrophobic amino acids from the amphipathic Nt17 segment, greatly affects its binding preferences and its tendency to localize to membranes of curvature.

**Figure 4:**
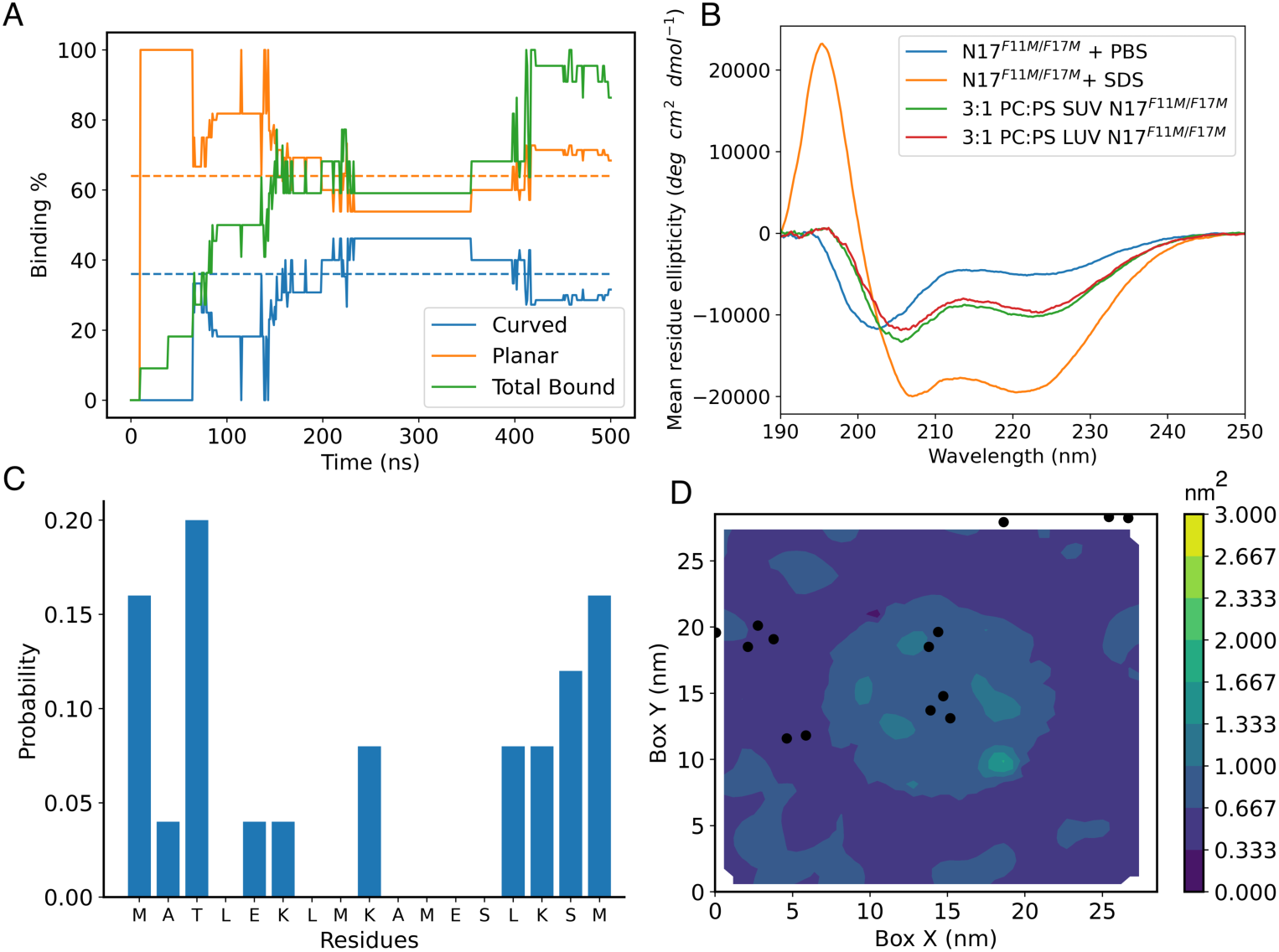
A mutated version of Nt17, Nt17 *^F^* ^11*M/F*^ ^17*M*^ has no specific binding preferences, binding to both the curved and planar regions. A) Binding percentages for one replicate of Nt17*^F^* ^11*M/F*^ ^17*M*^ to the curved and planar regions are denoted blue and purple respectively. The dotted lines represent the minimum percent binding required to qualify as having preference for curved and planar regions. The solid green line represents the percentage of membrane bound peptides in the system, B) Circular dichroism of Nt17*^F^* ^11*M/F*^ ^17*M*^ in varying solutions. 10 mM pH 7.3 PBS (blue); 10 mM pH 7.3 PBS with 2 mM sodium dodecyl sulfate (SDS) to induce helical folding (orange); 3:1 POPC:POPS SUVs in 10 mM pH 7.3 PBS (green); and 3:1 POPC:POPS LUVs in 10 mM pH 7.3 PBS (red). SUVs (small unilamellar vesicles) are around 30nm in diameter and LUVs (large unilamellar vesicles) are around 120 nm in diameter, C) First interaction probability for residues of Nt17*^F^* ^11*M/F*^ ^17*M*^ with the lipid headgroups, D) APL plot for one replicate of Nt17*^F^* ^11*M/F*^ ^17*M*^. Black dots indicate the center of mass of bound peptides.

Similar to Nt17, Nt17*^F^* ^11*M/F*^ ^17*M*^ also displays a high interaction probability at either termini, as seen in Fig. 4C. Since the mutation from F to M does not affect the amphipathic nature of Nt17*^F^* ^11*M/F*^ ^17*M*^, the mutated peptides also aggregate such that their hydrophobic face pointing inwards, resulting in first interactions that can occur from either the N or C terminal residues. Fig. 4D presents the APL for one replicate of the Nt17*^F^* ^11*M/F*^ ^17*M*^ system. Here, Nt17*^F^* ^11*M/F*^ ^17*M*^ binds to both M*_c_* and M*_p_*, displaying no preference for either region. Replicates of these simulations, shown in Figure S5, substantiate these trends.

### Mechanisms of curvature sensing

To better understand the mechanism of curvature sensing, we analyzed the effect of Nt17 binding on the membrane. Figure 5A illustrates the distribution of APL at M*_c_* for both the control simulation and the simulation with peptides. In the control, the APL at M*_c_* peaks around 1.5 nm^2^, with a median value of 0.8 nm^2^. In a planar POPC bilayer, our lipid model has APL values of 0.62 nm^2^,^30^ falling within the range of accepted values for POPC.^31,32^ The extension to 1.5 nm^2^ highlights the existence of packing defects, formed due to the curved region. However, these defects are enhanced in the presence of Nt17, where the APL distribution is vastly extended, ranging from values reaching up to 4 nm^2^, in sharp contrast to the condensed APL distribution of the control simulation. Figures 2 C,D tell the same story, with the curved region having higher APL values than the surrounding planar regions. In the control simulation, the curved region has APL values ranging from 0.625 nm^2^ to 1.562nm^2^, but upon binding of the peptide to the curved region, a jump in APL can be noted, with some regions having an APL of 2.5 nm^2^. Thus, binding of Nt17 further enhances lipid packing defects, corresponding to the extended tail in the APL distribution, seen in Figure 5A. The inset in Figure 5A presents a schematic of the process. Peptides may partition to different regions of the membrane surface either as a monomer, or as a small aggregate (such as a dimer/trimer/tetramer). Over time, the peptides bound to M*_c_* are able to diffuse towards each other, pushing the lipids apart and causing the increase in defects in the curved region of the membrane. Video S1 presents a short video of peptides interacting with the lipid bilayer. Note the peptides (each denoted by 2 red dots) first binding at different regions of the M*_c_*, then diffusing together, enhancing the packing defects and increasing the APL in the curved region.

**Figure 5:**
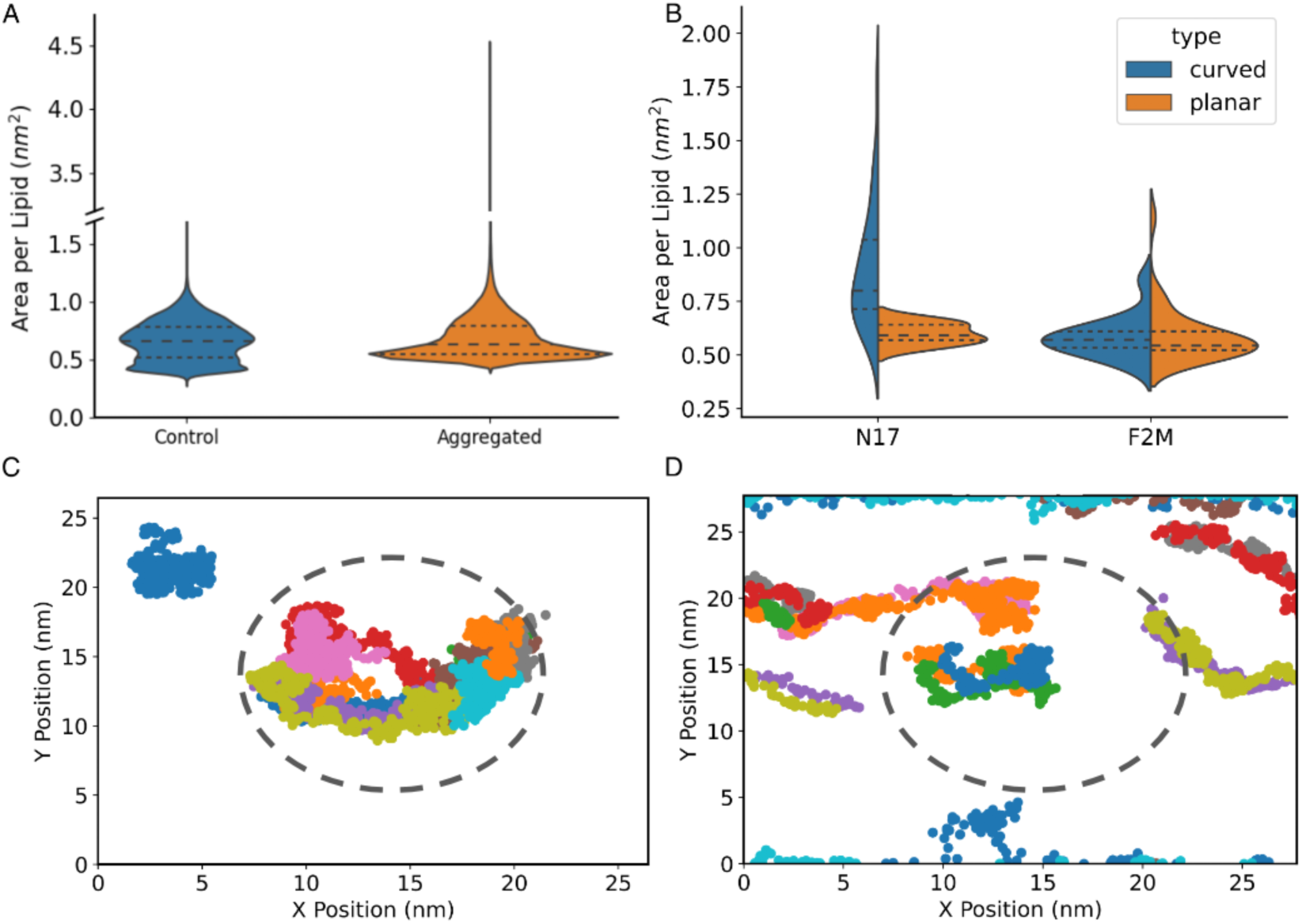
Nt17 detects regions of curvature by sensing lipid packing defects. A) Violin plots plot of area per lipid (APL) of the curved region of the membrane in the control simulation (no peptide) and simulation with Nt17. The inset contains a schematic describing the mechanism by which Nt17 can enhance membrane defects, ie. Nt17 binds to regions with packing defects and these defects consolidate into larger defects leading to the extended range of APL values, B) Area per lipid (APL) at the site of first peptide interaction for both Nt17 and Nt17*^F^* ^11*M/F*^ ^17*M*^, C) and D) present diffusion maps for Nt17 and Nt17*^F^* ^11*M/F*^ ^17*M*^ respectively. Positions denote the center of mass of the peptide and each peptide is marked in a different color. The dotted line roughly corresponds to the curved region of the membrane architecture.

We have confirmed that Nt17 is sensitive to membrane curvature through its affinity for lipid packing defects, and lack of the bulky hydrophobic groups in Nt17*^F^* ^11*M/F*^ ^17*M*^ results in loss of curvature specificity. To distinguish the binding of Nt17 and Nt17*^F^* ^11*M/F*^ ^17*M*^, we considered APL at the site of initial peptide binding, as seen in Fig. 5B. Due to its sensitivity for curved regions, Nt17 displays distinct partitioning preferences when interacting with M*_c_* and M*_p_* respectively. It is entropically more favorable for the large Phe residues of Nt17 to find regions of higher APL (and thus more space between headgroups) to bind. These regions primarily occur in the curved region of the membrane. In the case of Nt17*^F^* ^11*M/F*^ ^17*M*^, however, the distribution of APL at the initial peptide binding event are virtually identical when binding occurs at the curved or planar regions. The smaller size of methionine allows Nt17 to bind to regions where the defects are not nearly as large, as evidenced by the indiscriminate binding to both planar and curved regions. An interesting feature observed in the simulations of Nt17*^F^* ^11*M/F*^ ^17*M*^, is the diffusion of the peptide across regions of different curvatures, something not observed in the simulations with Nt17. In one replicate, we observed Nt17*^F^* ^11*M/F*^ ^17*M*^ initially binding to the planar region of the membrane, then diffuse to the curved region such that the binding percentage at the curved region is higher (see Figure S6B).

Figures 5C and D present diffusion maps for one replicate of Nt17 and Nt17*^F^* ^11*M/F*^ ^17*M*^, respectively. Here, the center of mass of peptides in direct contact with the membrane is tracked over time, and each peptide is marked in a different color. Nt17 and Nt17*^F^* ^11*M/F*^ ^17*M*^ demonstrate vastly different behaviour while interacting with the lipid. Nt17 binds to the curved region of the bilayer (center of simulation box), and displays very little diffusion. On the other hand, Nt17*^F^* ^11*M/F*^ ^17*M*^, diffuses over the entire membrane surface. We can connect back to Figure 5B, where we observed Nt17 binding to regions of high APL to accommodate its bulky hydrophobic groups, which then reduces the diffusion along the membrane surface. The smaller sidechains of Nt17*^F^* ^11*M/F*^ ^17*M*^ allow it to diffuse along the membrane, even from the curved region to the planar and vice-versa.

## Conclusions

In this work, we apply coarse-grain molecular dynamics simulations to investigate the interactions of Nt17, the N-terminal domain of htt protein, with curved membranes. A plethora of studies have detailed the importance of Nt17, not just in Huntington’s Disease pathology, but also in healthy cellular function. As the membrane binding domain of htt, it is crucial to understand how Nt17 interacts with the complex cellular environment, specifically, how Nt17 associates with curved membranes, a common intracellular feature.

The curvature sensing ability of Nt17 has been noted in prior experiments via AFM microscopy and EPR spectroscopy.^13,14^ We aimed to uncover a molecular mechanism that can explain this behavior. To do so, we use a membrane architecture containing both planar and curved regions, as depicted in Figure 1. The curved region (M*_c_*) accounts for 36% of the upper leaflet surface. Thus if more than 36% of the peptides are bound to M*_c_*, the peptide was classified as a curvature sensor.

Nearly 78% Nt17-M*_c_* binding was observed, firmly establishing Nt17 as a curvature sensor. We found that membrane binding is largely dependent on the C-terminal Phe residue, with over 30% of interactions occurring through this residue. Nt17 domains arrange as bundles in solution due to their amphipathic nature,^24^ thus interactions can occur through either termini. Since the C-terminus is more hydrophobic, we observe most interaction from this terminus. Mutation of the Phe residues to M, in Nt17*^F^* ^11*M/F*^ ^17*M*^, revealed the pivotal role that bulky hydrophobic residues play in curvature sensing. Nt17*^F^* ^11*M/F*^ ^17*M*^ displays no curvature binding preferences, interacting with both M*_c_* and M*_p_*.

These findings were found to be in agreement with circular dichroism spectra. Nt17 could differentiate between SUVs (more curvature) and LUVs (less curvature), as seen by the variance in signal intensity in Figure 2B. By contrast, Nt17*^F^* ^11*M/F*^ ^17*M*^ does not distinguish between LUVs and SUVs to the same extent. In the case of Nt17, the bulky hydrophobic groups gravitate to regions with higher APL, where it is entropically more favorable to bind. This occurs primarily in the curved region of the membrane, where lipid packing defects are abundant. Nt17*^F^* ^11*M/F*^ ^17*M*^ does not discriminate between curved and planar to the same extent, as shown by the similar APL distributions at the sight of first lipid interaction. Here, the smaller M residues are able bind to regions of the membrane, even if the area per lipid is low.

Two main strategies have been proposed to explain curvature sensing phenomena.^33,34^ Certain peptides, such as the BAR domain proteins, possess a slightly curved surface populated with charged residues, resulting in electrostatic attraction to curved membranes. On the other hand, ALPS motifs bind preferentially to curvature through sensing the lipid packing defects in curved membranes. They often possess large hydrophobic residues on one face and small polar groups (serine and threonine) on the polar face, leading to membrane curvature being a necessity for binding. α-synuclein is one such curvature sensor, with a poorly developed hydrophobic face, but a comparatively stronger polar face, granting it sensitivity to curved anionic lipid bilayers. Interestingly, mutating key polar residues to W, results in α-synuclein developing a sensitivity to zwitterionic lipid curvatures as well.^20^ Unlike α-synuclein and ALPS motifs, Nt17 has a well developed hydrophobic face, with 2 aromatic Phe residues and a sizable polar face of 5 charged residues. Thus, the mechanism by which Nt17 senses curvature is distinct. While Nt17 is able to detect and preferentially bind to membrane curvature, it can also bind to planar membranes, placing it in a unique category of curvature sensor. This dual binding behavior suggests that Nt17 could play a more nuanced role in cellular processes, potentially impacting how it facilitates membrane localization, aggregation, or signaling events within cells. Such flexibility in membrane interaction may influence downstream pathways in cellular trafficking, protein localization, or signal transduction.

The insights obtained from this study will be crucial to better develop our understanding of neuro-degenerative diseases, with Huntington’s Disease as a model system. A large body of work has highlighted the link between curvature sensing and curvature generation, with some peptides sensing curvature at low concentrations and generating curvature at high concentrations.^35,36^ Although our simulations employ high concentrations, we cannot assess the potential curvature generation due to the restraining potentials applied to retain the hemispherical-planar architecture. Our future works will feature updated lipid systems to examine possible membrane curvature generation upon Nt17 binding.

## Methods

Simulations were performed on GROMACS 2019.4 simulation engine.^37^ VMD 1.9.4^38^ was used for visualization. Analysis was performed on MDAnalysis 1.1.1^39^ using in-house scripts.

### Simulation parameters

#### Forcefield

In this work, we use the previously developed ProMPT forcefield^40^ to set up all simulations. ProMPT is a coarse-grained protein force-field that employs free-energy based parameterization and a 4:1 bead mapping. To account for the orientation of the atoms underlying the coarse grained beads, flexible dipoles consisting of opposing charges are incorporated into the coarse-grained beads. The electrostatic interactions induced by these dipoles introduce structural polarization to the model, allowing us to capture secondary structure changes in response to fluctuations in the local environment. ProMPT has been validated on a number of small proteins^41^ and more recently was employed to study aggregation of Nt17 and the following polyQ domain.^24^ Figure S8 presents the ProMPT representation of the protein. A helical dihedral potential with competing minima was applied to the Nt17 backbone beads, utilizing a force constant of 10 kcal/mol (see Figure S9). The lipid model, titled WEPMEM, is similarly designed, in that it features a 4:1 bead mapping with the addition of explicit dipole to polar coarse grained beads.^30^ The protein and lipid forcefields are used in conjunction with the polarizable Martini water.^42^

#### Curved membrane setup

Experimentally, curvature is typically studied either using vesicles or supported bilayers. To reproduce curvature in silico, membrane architecture containing both curved and planar regions was created using the BUMPy package.^43^ The hemispherical-planar membrane architecture pictured in Figure 1 features POPC, a phospholipid routinely encountered in the cellular environment and has a diameter of approximately 15 nm. To retain the shape of the membrane through the course of the simulation, a grid of dummy particles was restrained close to the phosphate group of the lipid, as detailed in the BUMPy paper.^43^ These dummy particles have a repulsive interaction with the acyl tail of the lipids (C_12_= 0.02581 kJ mol*^-^*^1^ nm^2^), ensuring that the membrane shape is maintained through the course of the simulation. The dummy particles do not engage in interactions with any other molecules in the system. To promote binding of Nt17 to the positive curvature (upper leaflet), we instituted a repulsive interaction (C_12_= 0.02581 kJ mol*^-^*^1^ nm^2^) between the headgroup of the lower leaflet with the protein backbone bead, ensures that peptides do not bind to the lower leaflet.

To stabilize the curved setup, an energy minimization of 5000 steps was performed, followed by an NVT equilibration for 10 nanoseconds with soft restraint on phosphate group. Next, an NPT simulation was run for 50 nanoseconds without any restraints on the phosphate. The final diameter of the membrane architecture is around 15 nm.

#### Simulation Setup

The control simulation (no peptide) was run for 250 ns using the NPT ensemble. For the simulations with Nt17 and Nt17*^F^* ^11*M/F*^ ^17*M*^, peptides were run added to the water box for a peptide concentration of approximately 20mM, amounting to 36 peptides and approximately 1400 POPC molecules, or peptide:lipid ratio of 1:40. The simulation box had x,y,z dimensions of 26nm, 26nm and 13 nm respectively. Following this, a short energy minimization of 5000 steps and equilibration of 5000 steps was performed. Counterions and a salt concentration amounting to 125mM were then added to the simulation box. 3 consecutive equilibrations of 5000 steps were performed with timesteps of 1fs, 5fs and 10fs respectively. Production was run for 500 ns with the NPT ensemble.

The Nose-Hoover thermostat, with a time constant of 1 picosecond, was utilized to maintain a fixed temperature of 300K and the Parrinello-Rahman barostat kept the pressure constant at 1 bar. A relative dielectric constant of 2.5 was applied, and electrostatic calculations were performed using Particle Mesh Ewald. Constraints were evaluated using the LINCS algorithm, and the neighbor list was updated every 10 steps.

### Analysis of simulations

To conduct the analysis of the simulations, we first established specific criteria. A distance cutoff of 7 Å was used to evaluate contacts across all analyses. Peptides were considered interacting or bound to another peptide, or adsorbed to the membrane, if they had five or more heavy atom contacts with the interacting partner (either another peptide or the lipid phosphate group). We further categorized peptide interactions with the lipid as either direct or indirect. Direct interactions were defined as a peptide having five or more heavy atom contacts with the phosphate group of the lipid. Indirect interactions were defined as instances where peptide X did not directly interact with the lipid bilayer but instead interacted with another peptide Y, which was in contact with the bilayer. In this scenario, peptide X is considered to be bound to the lipid indirectly through its interaction with peptide Y.

Analysis performed on the proteins considered all heavy atoms (backbone and sidechain beads) unless explicitly mentioned. For the lipids, only the phosphate bead was considered.

To calculate the percentage of peptides bound to the curved and planar regions shown in Figure 2A, we counted the number of direct and indirect peptides interacting with M*_c_* and M*_p_* using the above mentioned criteria, and normalized by the total number of bound peptides in the system at each time point. Total bound percentage is calculated as the sum of peptide directly and indirectly interacting with the membrane divided by the highest number of peptides bound.

Area per lipid plots shown in Figures 2 and 4 were calculated from the FATSLiM package,^44^ which provides the APL for each lipid in the system. Here, the black dots depict the center of mass of peptides bound to the curved region.

To classify the initial interaction between the peptide and lipid (Figures 3A and 4C), we defined the first interaction as lasting for at least 5 ns. This approach helped eliminate any transient interactions. The probabilities shown in the first interaction plots are averaged across all replicates, representing the frequency of peptide-lipid interaction per residue.

Figure 5 was created using the violinplot function from the seaborn package. Diffusion plots were created by plotting the center of mass of the peptides in direct contact with the membrane. Here, peptides are differentiated by color.

### Peptide preparation

Nt17, the first 17 amino acids of htt (MATLEKLMKAFESLKSF), and Nt17*^F^* ^11*M/F*^ ^17*M*^ (Nt17 with an F11M/F17M substitution), both with C-terminal amidation, were obtained via custom synthesis (GenScript, Piscataway, New Jersey). Disaggregation and solubilization of Nt17 and Nt17*^F^* ^11*M/F*^ ^17*M*^ peptides was achieved using established protocols.^45^ Briefly, peptide was dissolved overnight at a concentration of 0.5 mg/mL in a 1:1 mixture of trifluoroacetic acid (TFA, Sigma) and hexafluoroisopropanol (HFIP, Sigma). After vortexing, a N_2_ stream was used to evaporate solvent. To remove residual solvent, samples were placed in a vacufuge concentrator overnight, resulting in thin peptide films which were then stored at *-*20*°*C. For each experiment, peptide films were resuspended in ultrapure water that had been adjusted to pH 3 with TFA and then diluted in phosphate-buffered saline (PBS) (10 mM phosphate, 138 mM NaCl, 2.7 mM KCl, pH 7.3) (Dulbecco) to a suitable final concentration, typically 100-200 µM.

### Vesicle preparation

Large unilamellar vesicles (LUVs) and small unilamellar vesicles (SUVs) composed of 3:1 1palmitoyl-2-oleoyl-glycero-3-phosphocholine (POPC): 1-palmitoyl-2-oleoyl-sn-glycero-3-phosphoL-serine (POPS) (Avanti Polar Lipids, Alabaster, AL) were prepared using extrusion and sonication methods. Briefly, a thin lipid film of lipid, dried under nitrogen from a chloroform stock, was hydrated at 2.5 mg/ml in PBS (10 mM phosphate, 138 mM NaCl, 2.7 mM KCl, pH 7.3) and vortexed for 10 min. For LUVs, this solution was treated with 10 freeze-thaw cycles using ethanol/dry ice and warm water baths and then extruded 11 times through a 100 nm pore polycarbonate filter (Avanti Polar Lipids). For SUVs, this solution was bath sonicated for 30-60 min. Dynamic light scattering measurements with a Zetasizer ZS90(Malvern, Worcestershire, UK) were used to measure vesicle diameter (LUVs 130 nm; SUVs 30 nm) and ensure sample homogeneity.

### Circular dichroism

CD spectra were collected on a JASCO J-1500 spectropolarimeter using a 1 mm pathlength quartz cuvette. Samples contained 0.15 mg/mL peptide (76 µM) in PBS at pH 7.3 and spectra were normalized to protein concentration measured by UV spectroscopy at 205 nm (*є_Nt_*_17_= 65340 M*^-^*^1^ cm*^-^*^1^; *є_Nt_*_17_*F* 11*M/F* 17*M* = 58570 M*^-^*^1^ cm*^-^*^1^).^46^ Lipid interfaces were introduced via 1.25 mg/mL lipid (1.60 mM) SUVs or LUVs. Spectra are baseline corrected and averaged over three scans, measured from 260 nm to 190 nm with a 0.2 nm data interval, and 4 s data averaging.

## Acknowledgement

This research was supported by the National Science Foundation under grants CHE-1454948, CHE-2202281 and a Major Research Instrumentation grant 1725534. The authors acknowledge the University of Maryland supercomputing resources (http://hpcc.umd.edu) made available for conducting the research reported in this paper.

## Supporting Information Available

Binding fraction for replicates of Nt17 and Nt17*^F^* ^11*M/F*^ ^17*M*^ simulations, average aggregate sizes for Nt17 and Nt17*^F^* ^11*M/F*^ ^17*M*^, APL plots for Nt17 and Nt17*^F^* ^11*M/F*^ ^17*M*^, comparison of binding interactions between Nt17 and Nt17*^F^* ^11*M/F*^ ^17*M*^, peptide diffusion plots for both Nt17 and Nt17*^F^* ^11*M/F*^ ^17*M*^, coarse-grained representation of Nt17 and the dihedral potential applied to the backbone beads of Nt17 and Nt17*^F^* ^11*M/F*^ ^17*M*^, video presenting the enhancement of lipid packing defects upon binding of Nt17 binding to the membrane.

## Notes

### Competing Interest Statement

The authors have declared no competing interest.

### Summary of Updates

This version features updates to percentage of binding and diffusion plots and revised explanations.

