## Supplementary Information for "Investigating curvature sensing by the Nt17 domain of Huntingtin protein"

### Supplelementary Information

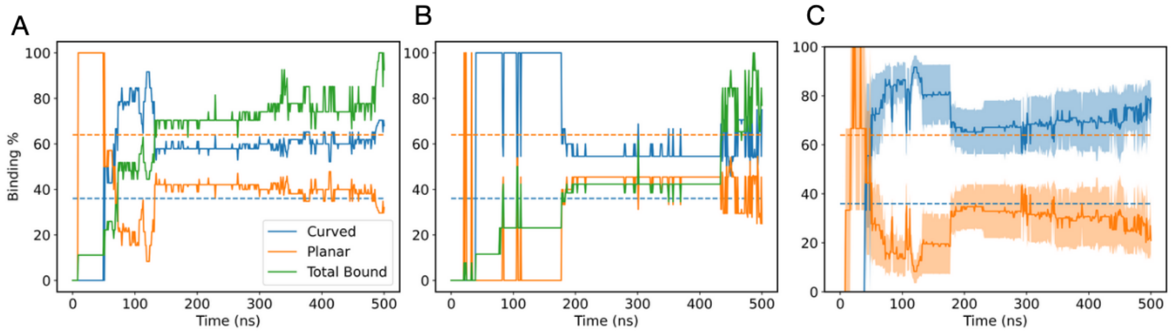


Figure S1: A, B) Percent binding for 2 other replicates of Nt17 with the planar-hemispherical membrane architecture. C) Percent binding of calculated across the 3 replicates. The solid line represents mean value and shade indicates the standard error across replicates. Dotted lines represent the percentage required to qualify preference for the curved and planar regions respectively.

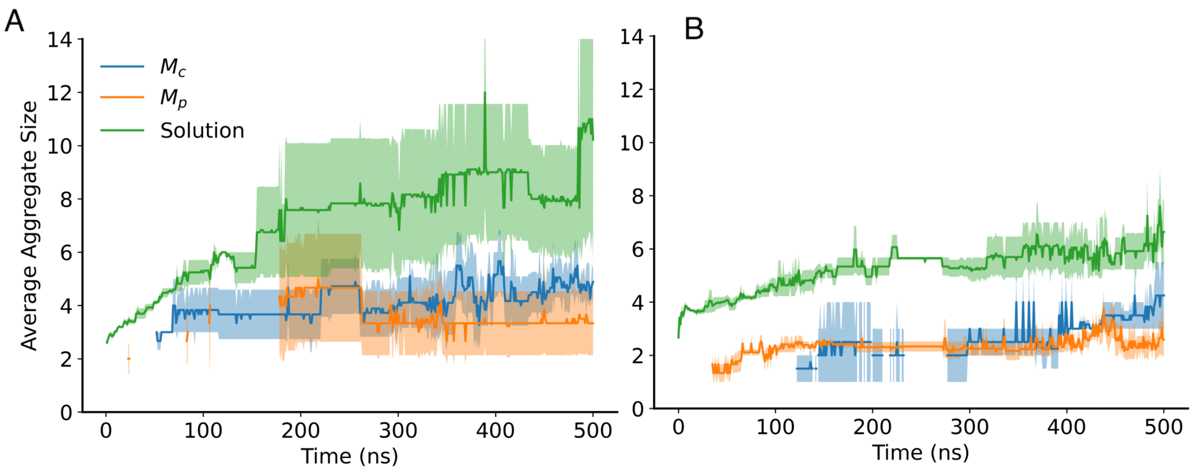


Figure S2: Average aggregate size for A) Nt17 and B) Nt17$^{F11M/F17M}$ on the curved region($M_c$), planar region($M_p$) and in solution. The solid line is the mean value and shade indicates the standard error across replicates.

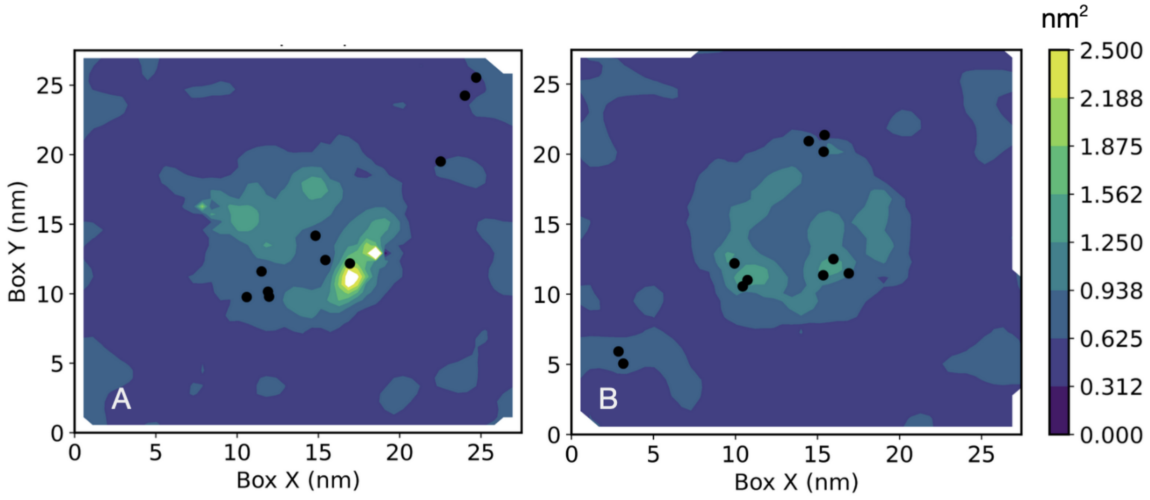


Figure S3: Area per lipid plots for the replicates of Nt17 simulation. Black dots indicate the center of mass of peptides that have bound to the membrane.

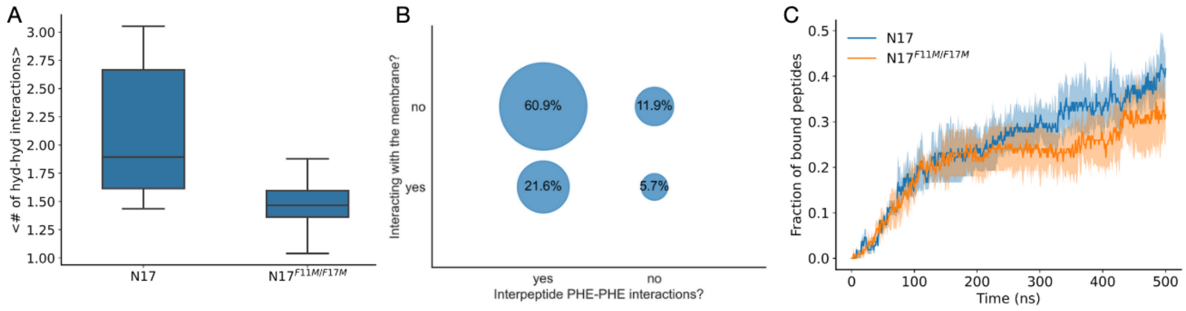


Figure S4: Nt17 and Nt17$^{F11M/F17M}$ have similar cumulative binding fraction, but vary in the size of the hydrophobic core formed in the oligomeric state. A) Average number of hydrophobic sidechain interactions experienced by Nt17 oligomers in solution. B) Binary classification of competing interactions experienced by Nt17- either engaged in hydrophobic interactions with other peptides, or the F sidechains are interacting with the membrane. C) Fraction of Nt17 and Nt17$^{F11M/F17M}$ bound to the membrane.

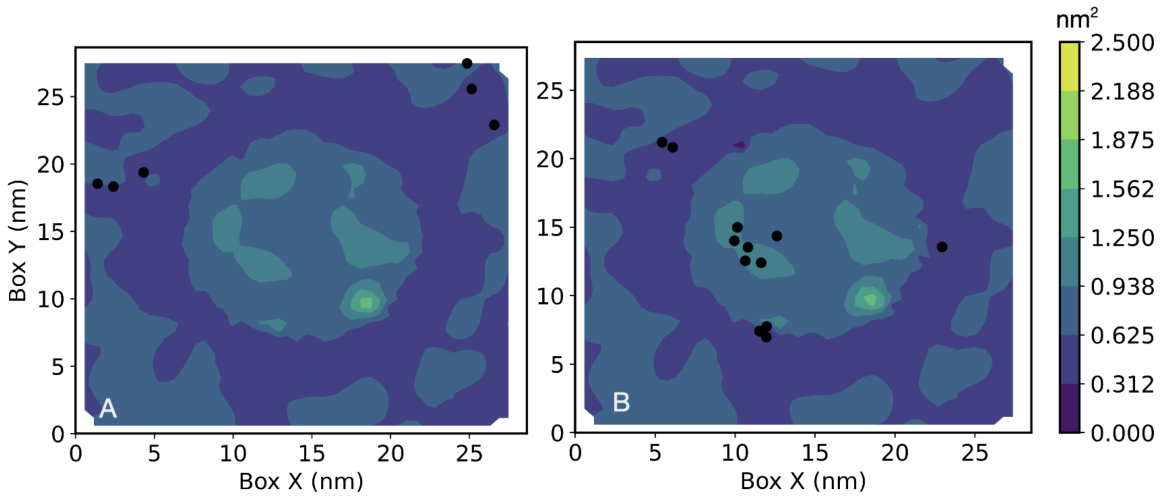


Figure S5: Area per lipid plots for the replicates of Nt17$^{F11M/F17M}$ simulation. Black dots indicate the center of mass of peptides that have bound to the membrane.

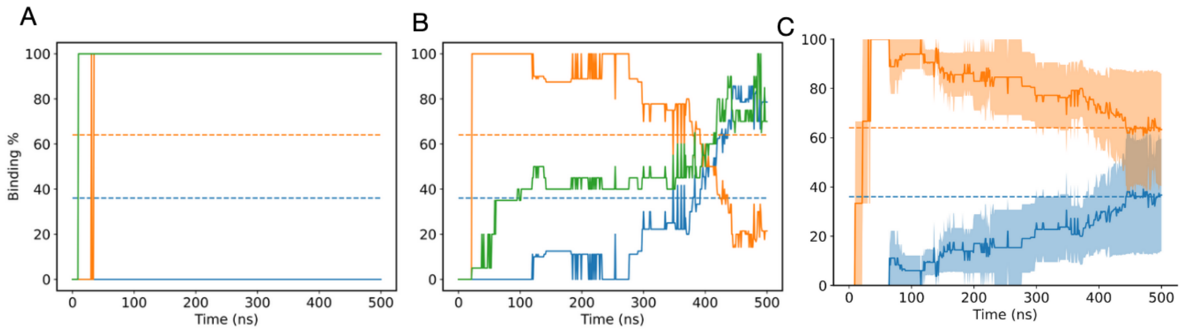


Figure S6: A, B) Percent binding for 2 other replicates of Nt17$^{F11M/F17M}$ with the planar-hemispherical membrane architecture. C) Percent binding of calculated across the 3 replicates. The solid line represents mean value and shade indicates the standard error across replicates. Dotted lines represent the percentage required to qualify preference for the curved and planar regions respectively.

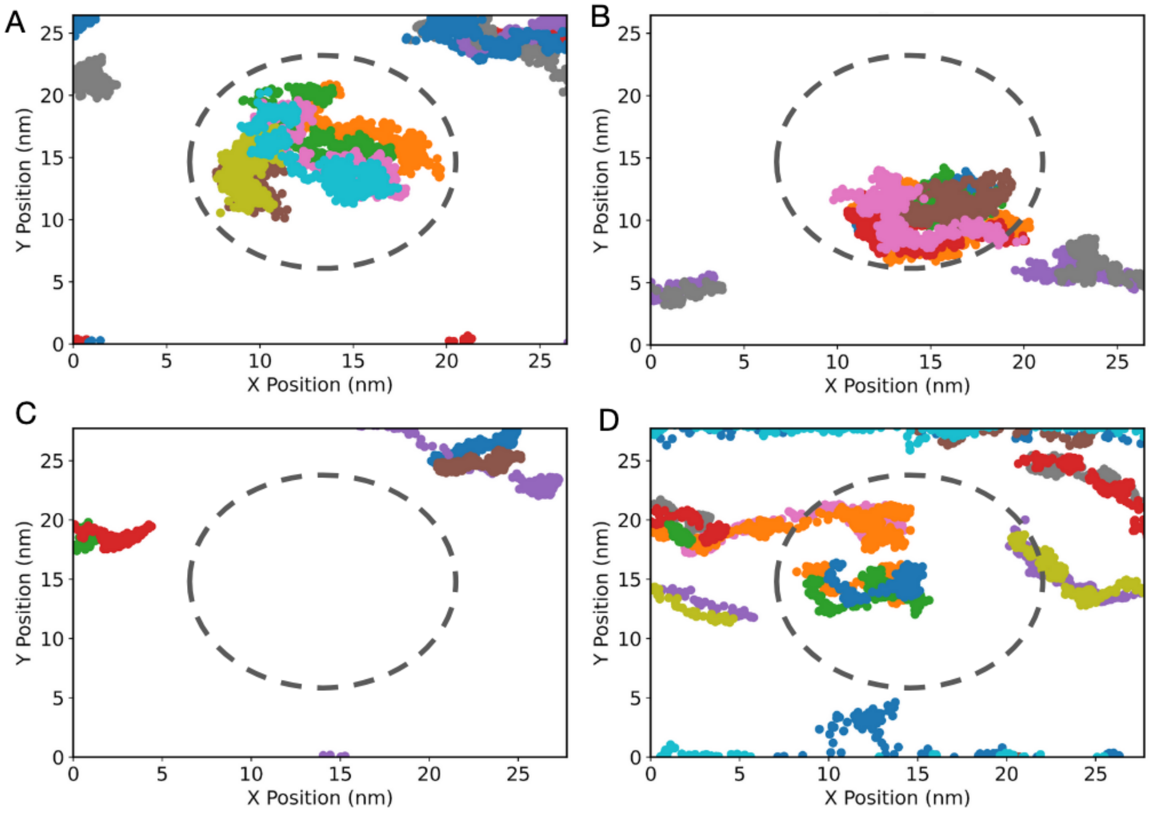


Figure S7: Diffusion maps created for A,B) replicates of Nt17 and C,D) replicates of Nt17$^{F11M/F17M}$. Center of mass of each interacting peptide over the course of the simulation is demarcated in a different color. The dotted line roughly corresponds to the curved region of the membrane architecture.

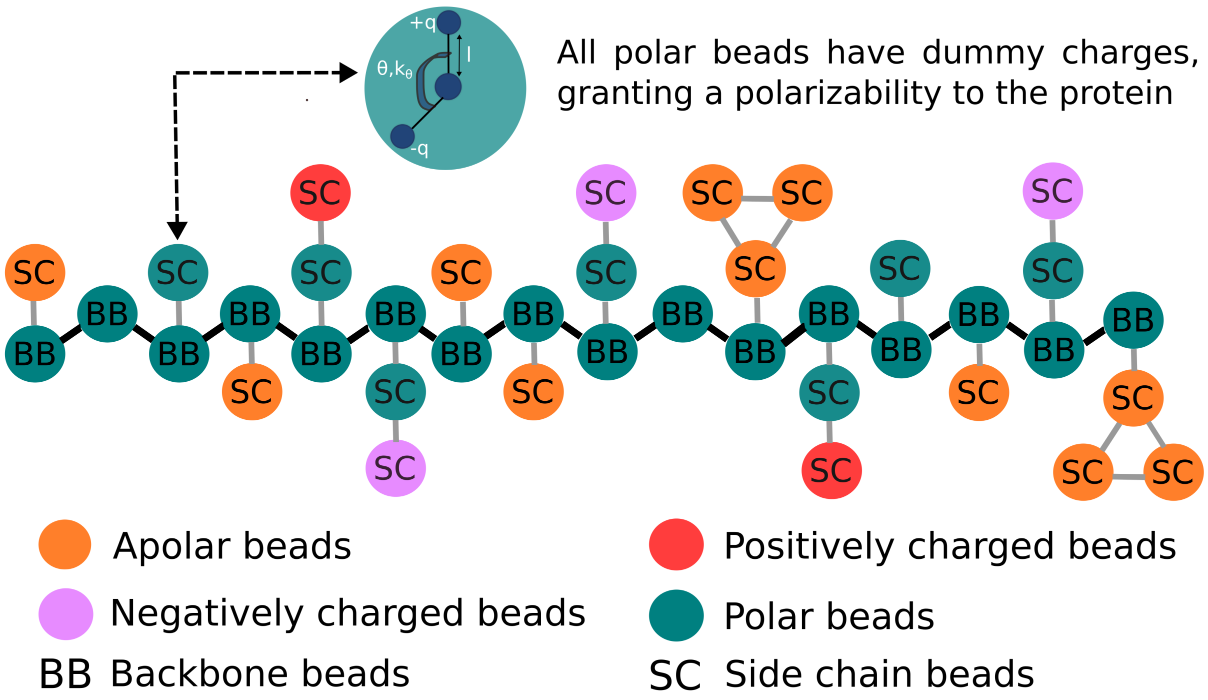


Figure S8: Coarse grained representation of the Nt17 domain. The forcefield was implemented as described in the ProMPT paper,[1] with the exception of cation pi interactions.

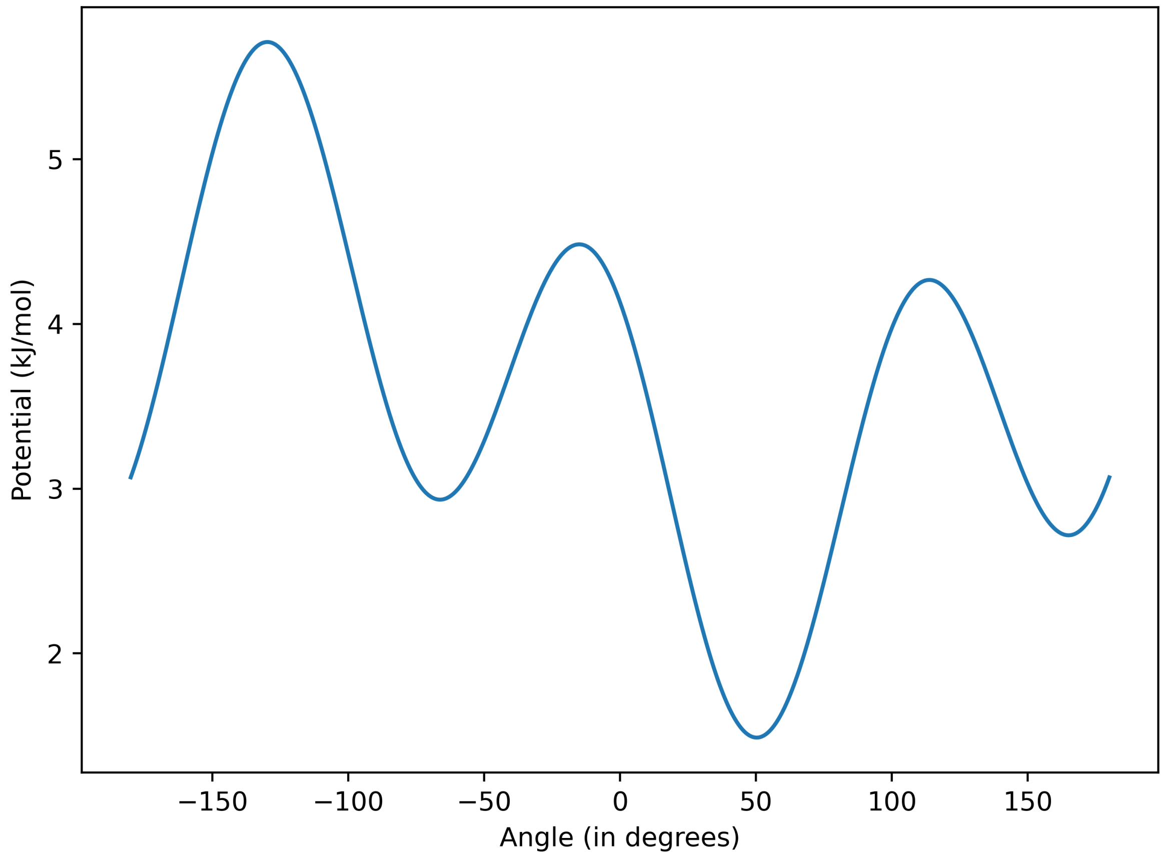


Figure S9: Dihedral potential applied to the backbone beads of the Nt17 domain.
